# Mechanism of MEK1 activation by phosphorylation

**DOI:** 10.1101/2025.10.01.679899

**Authors:** Everett Jin, Levent Sari, Milo M. Lin

## Abstract

Phosphorylation is the most common post-translational protein modification, often acting as the activator of protein function. In the case of MEK1, a member of the MAP kinase family, phosphorylation of both S218 and S222 are necessary for activation. Yet the molecular activation mechanism and its cooperative nature are poorly understood, especially due to the lack of experimental phosphorylated MEK1 structures. We performed molecular dynamics simulations to investigate the structural and dynamical consequences of single and double phosphorylation in MEK1. We find that successive phosphorylation progressively unwinds the helix containing the phosphorylation sites, thereby rotating the phosphate groups to directly interact with the catalytic site. Consequently, the solvent accessible surface area of the catalytic residues increase with phosphorylation. Yet, only in the double-phosphorylated state do all four critical catalytic residues become solvent exposed. By calculating the conformational entropy, we find that only in the double-phosphorylated state do all four catalytic residues have mutual information with each other, suggesting that phosphorylation also induces correlated motion at the active site. To validate our approach, we show that our simulations predict the alignment of the R-spine motif in the double-phosphorylated state, in agreement with the other kinases for which this alignment has been experimentally observed. These results show that phosphorylation can, via a partial unfolding mechanism, increase the solvent exposure of and dynamical coupling within the active site, both of which are critical for enzymatic activity.

## Introduction

Protein kinases play a vital role in the cell, including controlling apoptosis, cell division, metabolism, and DNA replication, mainly as part of regulating diverse signal transduction pathways (*1*) (*2*) (*3*) (*4*) (*5*). Between thirty and fifty percent of proteins are substrates of kinases, including many kinases themselves (*2*) (*6*). As a result of their significant role in signaling, misregulation or malfunction of protein kinases is present in many diseases. For example, many cancer-targeting drugs are kinase inhibitors (*1*) (*7*) (*8*). Kinases contain two conserved lobes, a small C-lobe primarily associated with ATP-binding, and a larger N-lobe associated with catalysis and peptide binding (*9, 10*). Between these two lobes lies a cleft containing the active site, which houses the ATP molecule that serves as the phosphate donor to the phosphorylation substrate (*10*) (*11*). Near the opening of the cleft lies the substrate binding site (*11*). Most kinases are themselves activated by phosphorylation at specific sites (*1*). Many such kinases are activated by phosphorylation of a residue or residues on a surface activation loop near the active site (*12*) (*13*). For example, in protein kinase A (PKA), a single phosphorylation of threonine turns the protein from an inactive to an active state (*14*). On the other hand, the activity of mitogen- and stress-activated kinase 1 (MSK1) is regulated by many phosphorylation sites (*15*).

A comparison of serine/threonine and tyrosine kinases showed that when these kinases are phosphorylated, a conserved three-residue DFG kinase motif flips from the “DFG-out” conformation to the “DFG-in” position. The proposed activation mechanism suggests that this flip facilitates stabilization of the kinase in its activated conformation (*16*). In PKA, near the DFG motif, the catalytic loop becomes more solvent accessible when the kinase is phosphorylated, although the structural and dynamical changes that cause this change were not resolved (*14*). A second structural response to phosphorylation is the formation of two hydrophobic spines: a regulatory spine which forms salt bridges and hydrogen bonds with the rest of the kinase, and a catalytic spine which aids in ATP localization (*16*) (*17*). In protein kinase A (PKA), it was found that the phosphoryl group added as a result of phosphorylation facilitated additional hydrogen bonding, including in regions around the active site of the protein (*18*).

The dynamics of certain protein kinases in response to phosphorylation have also been studied through molecular dynamics simulations for enzymes like JNK3, WNK, and BAK1 (*19*) (*20*) (*21*). In c-Jun N-terminal kinase 3 (JNK3), for example, both the *α*C helix in the N-lobe and the activation loop switch from a “closed” to an “open” conformation when JNK3 is phosphorylated (*20*). However, the detailed mechanism relating such dynamical changes to kinase activation, including identification of changes in the accessibility and motion of active site residues, is poorly understood.

In this study, we used MEK1 as a model system to examine protein activation by phosphorylation (1). MEK1 is a MAP kinase, part of the well-studied MAPK pathways, which plays a role in major cellular processes, including proliferation, differentiation, and apoptosis (*22*) (*23*). MEK1 is activated by the phosphorylation of two serine residues in the activation loop, Serine-218 and Serine-222, a highly conserved mechanism within MAP kinases (*24*). Both sites must be phosphorylated for MEK1 to be activated (*24*), yet the mechanism behind this cooperativity has not been resolved. In particular, the crystal structure of unphosphorylated MEK1 reveals that the DFG motif lies in a “DFG-in” conformation in which the aspartate within the motif is attracted to ANP. Therefore, in contrast to other kinases such as PKA, flipping of the DFG motif does not contribute to the activation mechanism of MEK1.

We performed all-atom molecular dynamics simulations on four ATP-bound MEK1 states: unphosphorylated, S218-phosphorylated, S222-phosphorylated, and double-phosphorylated MEK1. We show that, contrary to other kinases, MEK1 phosphorylation does not proceed via a DFG-flipping mechanism. Instead, our results showed that a partial unfolding of the activation loop alpha-helix. This partial unfolding allows S218 and S222 phosphate groups to directly interact with the active site, facilitating an increase in the catalytic residue solvent accessibility. We show that only in this double-interaction state do all four key catalytic residues become solvent exposed. We also demonstrate, using conformational entropy analysis, that only in the completely activated state does MEK1 have correlated movements within the four catalytic residues and in the region surrounding the active site, suggesting that the movement of residues in MEK1, in addition to its structure, is involved in its activation.

## Results

The structural components of MEK1 are shown in Fig. 1. The N-lobe includes a large beta-sheet, while the C-lobe is mainly alpha-helical. Like other kinases, the active site of MEK1 is located between the N lobe and the C lobe (*25*). The MEK1 catalytic loop consists of H188, R189, D190, and K192, the key residues that participate in the catalytic activity of MEK1 and transfer the phosphoryl group from ATP to ERK2 (*25*). A key residue in the catalytic loop is aspartate-190, which is believed to deprotonate ERK1 and facilitate its nucleophilic attack on ATP (*26*). In addition, K192, a conserved lysine in kinases, was found in PKA to be involved in the transfer of the phosphoryl group from ATP to the substrate rather than the anchoring of ATP (*27*).

**Figure 1:**
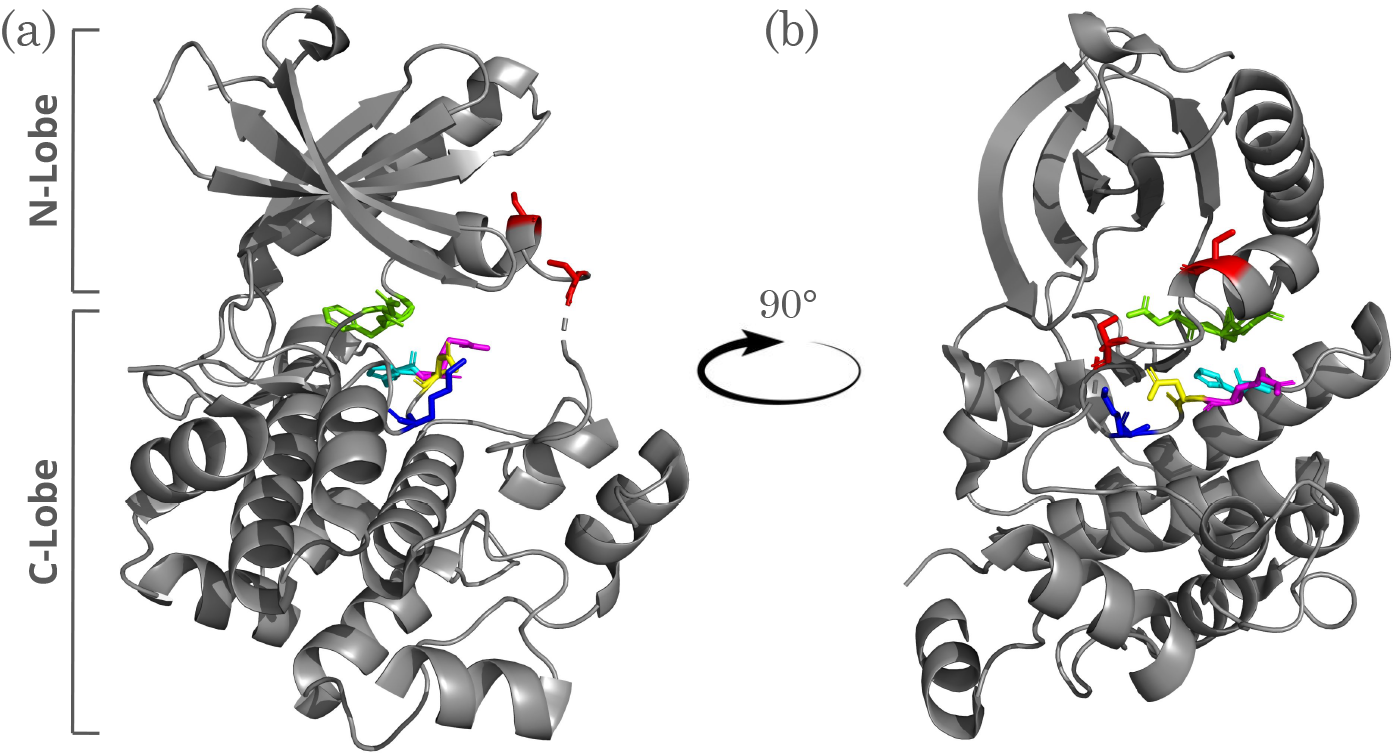
MEK1 crystal structure in the unphosphorylated state with ligands removed. (a) shows a front view, while (b) shows a side view. The critical catalytic residues are shown in stick format, with cyan indicating H188, magenta indicating R189, yellow indicating D190, and blue indicating K192. Red residues indicate phosphorylation sites. The DFG motif (residues 208-210) is highlighted in green. Since D208 is pointed towards the active site, the DFG motif is considered to be in the “in” conformation.

### Phosphorylated MEK1 Accesses an Activated, Solvent-Accessible State

Because solvent accessibility of the active site pocket is a criterion for kinase activity, we computed the total solvent-accessible surface area (SASA) of the four key catalytic residues of the active site in all four states for each system. We randomly selected a frame in which total active-site SASA is at the top one percent level as representative structures that are most competent for substrate binding (Figure 2). In both the unphosphorylated (Figure 2a) and S218-phosphorylated states (Figure 2b), the MEK1 activation loop adopts an alpha-helical conformation and is highly ordered throughout nearly all of the simulation times. In the unphosphorylated state of MEK1, the alpha-helical activation loop of MEK1 has three turns, with both S218 and S222 facing away from the active site (Figure 2a). In S218-phosphorylated MEK1, the alpha-helical structure of the activation loop, still with three turns, largely remains the same (Figure 2b). Although S218 is situated in the middle of the alpha helix, phosphorylation of S218 does not disrupt this helix because its sidechain faces away from the center of the helix. In addition, the residues around S218, which remain alpha-helical, keep S218 locked in place, like in the unphosphorylated state. A consequence of the relative rigidity of the activation loop, even when S218 is phosphorylated, is that S218 and S222 are not in proximity to the active site residues and therefore do not interact with them.

**Figure 2:**
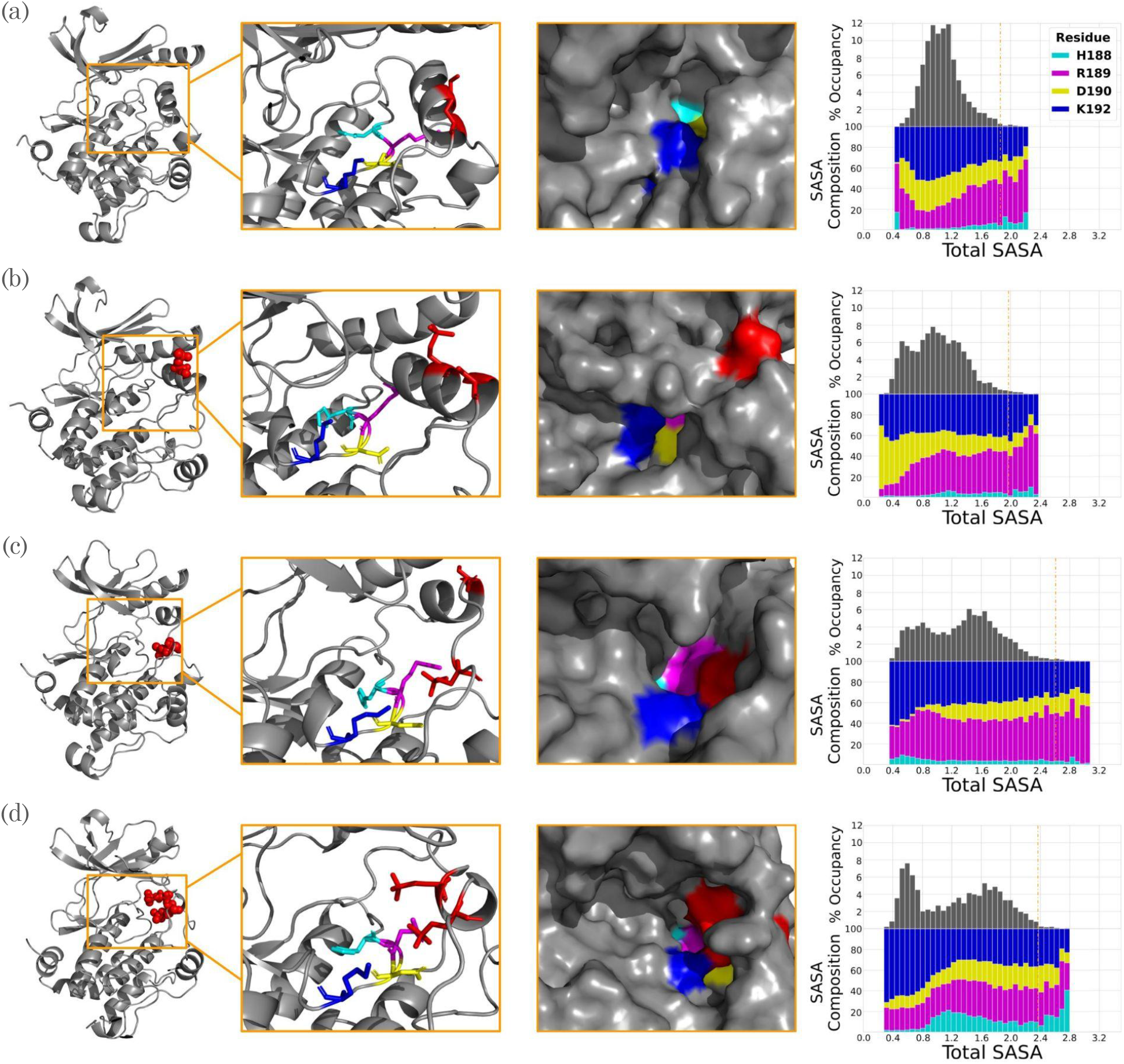
MEK1 structures at the one-percent occupancy level for the total SASA of critical active site residues. (a), (b), (c), and (d) represent unphosphorylated, S218-phosphorylated, S222-phosphorylated, and double phosphorylated MEK1, respectively. The panels show both a cartoon and a surface representation of the active site. The histograms show the SASA distribution of each MEK1 state. In both the panels and the histograms, cyan indicates H188, magenta indicates R189, yellow indicates D190, and blue indicates K192. Red residues indicate phosphorylation sites.

However, when S222 is phosphorylated, the alpha-helix becomes significantly more disordered (Figure 2c). Note that S222 is at the end of the alpha helix and forms the “bridge” between the alpha-helical part of the activation loop and the disordered region. When S222 is phosphorylated, its sidechain becomes attracted to that of R189, one of the critical active site residues. Because S222 is not contained within the alpha helix, it is able to orient its sidechain towards the active site. This causes a disordering of the alpha-helix, which is opposite to the stabilizing effect of phosphorylation on the activation loop previously found for other kinases (*19*) (*21*). However, S218 remains a part of an alpha-helical motif in the activation loop, and does not interact with the active site. Rather, it is at the edge of a two-turn alpha helix, positionally similar to S222 in unphosphorylated or S218-phosphorylated MEK1.

When S218 and S222 are both phosphorylated (Figure 2d), the alpha-helix disorders even more, as both S218 and S222 become attracted to R189. Since the alpha helix has already been disordered by S222’s attraction to R189, S218 can also move from its original position and orient its sidechain towards R189. This results in the alpha-helix motif within the activation loop having a single turn, located between S218 and S222. As a consequence, in the double-phosphorylated state, the side chains of S218 and S222 directly interact with active site residues, as shown in the surface renderings in Figure 2. To our knowledge, this is the first example of kinase activation through activation loop helix disordering, with particular importance for cooperative activation by multi-site phosphorylation.

To quantify the effect of phosphorylation on the active site structure, we calculated the SASA distribution of each of the individual catalytic residues for all four phosphorylation states. In the unphosphorylated and S218-phosphorylated states (Figure 2 (a,b)), the SASA distribution has a single peak at low SASA. Although the SASA distribution of S218-phosphorylated MEK1 has a larger standard deviation than that of unphosphorylated MEK1, the peak values of the distribution are at a similar total SASA, indicating that, for a majority of the trajectory, the critical catalytic residues of the MEK1 remain in a low-SASA conformation. In contrast, the total SASA distributions of S222-phosphorylated and double-phosphorylated MEK1 have two distinct peaks (Figure 2 (c,d)), suggesting that, in these two phosphorylation states, MEK1 is accessing two distinct conformational populations. The difference between S222-phosphorylated and double-phosphorylated MEK1 lies in the solvent accessibility of individual catalytic residues. In S222-phosphorylated MEK1, the total SASA of the catalytic residues is predominantly contributed by K192 and R189, as shown in the histogram in Figure 2 (c). The other catalytic residues, D190 and H188, either have very little SASA compared to K192 and R189, or little to no significant SASA at all. On the other hand, double-phosphorylated MEK1 has roughly an equal distribution of individual SASAs. The cyan part of the histogram, representing H188, covers a significantly larger portion of the total SASA than in S222-phosphorylated MEK1. The difference between the solvent-accessibility of individual catalytic residues suggests that although S222-phosphorylated MEK1 has a relatively large total SASA, single phosphorylation is not enough to solvent expose all critical catalytic residues simultaneously. Therefore, even though double-phosphorylated MEK1 has a comparable total active-site SASA to S222-phosphorylated MEK1, only in the double-phosphorylated case do all four catalytic residues, in particular H188, become solvent-exposed. These results therefore serve as a mechanistic explanation for the observed cooperativity of MEK1 activation.

### Conformational fluctuations of MEK1 catalytic residues become correlated in the double-phosphorylated state

To assess the dynamics of the critical catalytic residues, we used our in-house developed ciMIST framework (*28*) to calculate the conformational entropy and mutual information between residues in all four phosphorylation states, based on fluctuations of all dihedral and bond angles sampled in the MD simulations (Figure 3). In the unphosphorylated state, R189 has significant mutual information with seven residues, including D190. However, K192 and H188 do not have significant mutual information with any residues, suggesting that they do not move in coordination with R189 and D190, which inhibits catalytic activity in unphosphorylated MEK1. In the S218-phosphorylated state, R189 has significant mutual information with nine residues, including both D190 and H188. However, like the unphosphorylated state, K192 does not have any mutual information with any of the other residues, again consistent with the fact that experimentally, S218-phosphorylated MEK1 shows an insignificant increase in kinase activity (*24*). In the S222-phosphorylated state, there is very little significant mutual information between the critical catalytic residues and the rest of the protein. Like both S218-phosphorylated and unphosphorylated MEK1, K192 has no mutual information with the other critical catalytic residues in the S222-phosphorylated state.

**Figure 3:**
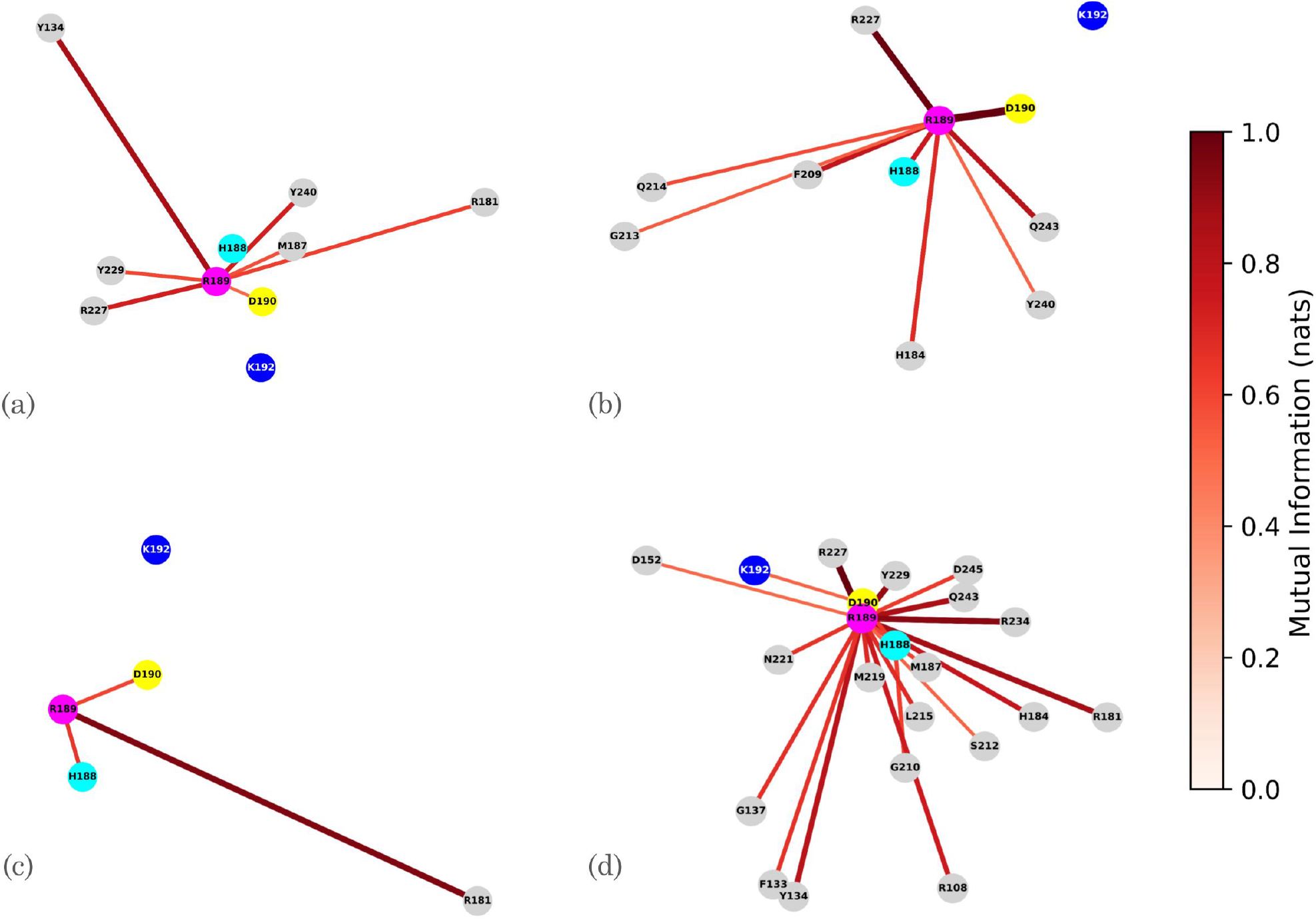
Mutual information network between the four catalytic residues and all other residues in (a) unphosphorylated, (b) S218-phosphorylated, (c) S222-phosphorylated, and (d) double phosphorylated MEK1. Edges represent any mutual information with the four critical catalytic residues exceeding 0.5 nats. Cyan represents H188, magenta represents R189, yellow represents D190, and blue represents K192. The Euclidean distances between the residues in the graphs are proportional to their distances in the MEK1 structures.

In contrast to the other three states, all four of the critical catalytic residues in double-phosphorylated MEK1 have significant mutual information with each other. This suggests that only in the double-phosphorylated state do all four critical catalytic residues have coordinated movements, which may be necessary for catalytic activity. While Figure 2 suggests that the solvent accessibility of H188 is crucial to catalytic activity, Figure 3 further suggests that the coordinated movements of the key catalytic residues is also essential. Specifically, in all cases other than the double-phosphorylated one, K192 does not have coordinated movements with the rest of the catalytic residues. Because K192 is a conserved lysine crucial for the phosphotransfer reaction of kinases, it may serve as a “gatekeeping residue” to catalytic activity (*27*). In fact, among the active-site residues, K192 is the only residue that directly interacts with the bound ATP, through the phosphate oxygen. These results show that, in the unphosphorylated or singly-phosphorylated states of MEK1, in addition to the structural barriers to activation presented by the solvent accessibility of critical catalytic residues, there exist signatures of cooperative active-site motion in the double-phosphorylated state.

To assess the influence of neighboring residues on the SASA of MEK1’s critical catalytic residues, for each state of MEK1, we determined the top 40 residues with the closest distance to the active site. We then collected the residues that were within the top 40 in all four MEK1 states. There were 35 such residues, including the four critical catalytic ones. Next, for each of the 35 residues identified, we employed the DBSCAN algorithm to cluster the MEK1 trajectory into different structural states based on the position of the residue in question (*29*). Then, we binned the total SASA of the critical catalytic residues, and calculated the mutual information between the position of the specific residue with the total SASA of the critical catalytic residues. As MEK1 progresses from unphosphorylated, S218-phosphorylated, S222-phosphorylated, to double-phosphorylated, the mutual information of each critical catalytic residue with SASA increases monotonically (Figure S1). Furthermore, the mutual information with SASA among the residues shown in Figure S1, which are those that are consistently close to the active site, increases in general as MEK1 becomes more activated. This suggests that, in addition to the active site becoming more coordinated, the residues of MEK1 surrounding the active site become more coordinated as well. In other words, the whole active site region accesses a less random, more coordinated state that allows MEK1 to perform its catalytic function.

### MEK1 phosphorylation leads to formation of the R-spine, a signature of kinase activation

Experimental structural characterization kinases at different levels of phosphorylation indicates that the alignment of four residues into an “R-spine” is a general feature of kinase activation (*18*) (*30*). The R-spine residues bridge both the N-lobe and C-lobe, and come from structurally and/or catalytically important parts of the kinase (*30*). Consistent with this picture, starting from the experimental structure of MEK1 in the unphosphorylated state in which the R-spine is not assembled (Figure 4a), our simulations predict the R-spine assembling as a consequence of phosphorylation (Figure 4b). This assembly is similar to that observed experimentally in proteins like PKA, for which the unphosphorylated (Figure 4c) and phosphorylated structures (Figure 4 (a) (d)) were experimentally determined (*18*).

**Figure 4:**
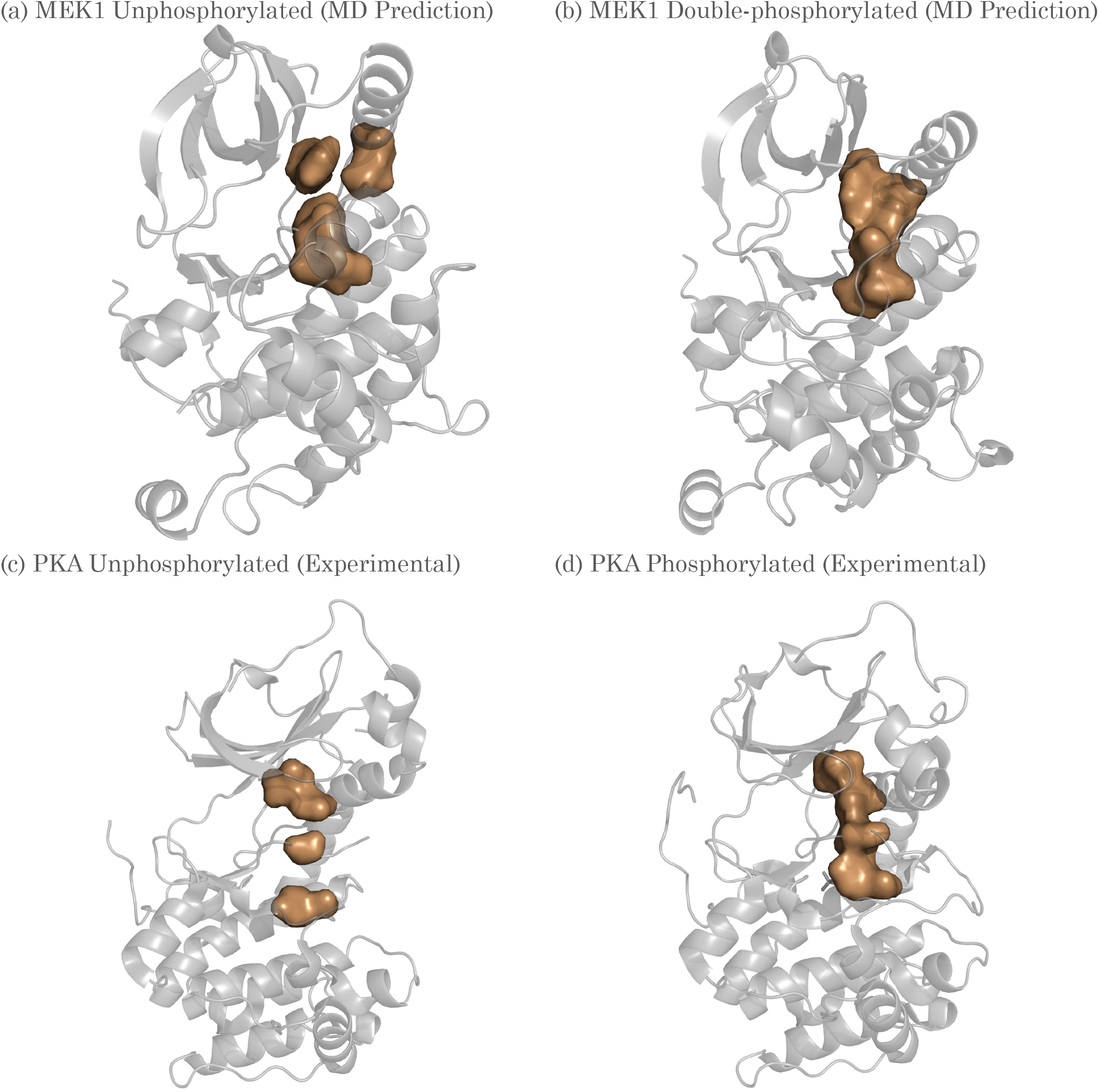
The R-Spine assembly of MEK1. (a) shows the R-spine in the centroid unphosphorylated structure of MEK1, and (b) shows the R-spine in the centroid double-phosphorylated, activated structure of MEK1. (c) and (d) are experimental PDB structures 1j3h and 4dfy showing the R-Spines in the inactive (unphosphorylated) and active (phosphorylated) state of PKA (*18*)

## Discussion

In this study, we used MEK1 Kinase as a model protein to study the effects of phosphorylation, one of the most common post-translational modifications, on kinase activation. Using all-atom molecular dynamics simulations, totaling 12 microseconds in total simulation time, as well as structural and dynamical analyses, we find that there are four main steps governing activation of kinases by phosphorylation: firstly, an unwinding of the activation loop alpha-helix, allowing the phosphorylation sites to interact with the active site; secondly, a roughly equally-distributed high SASA of critical catalytic residues; thirdly, the coordinated movements of all the critical catalytic residues; and lastly, the coordinated movements of residues around the active site, to form a kinase that can perform its catalytic function.

Because phosphorylation causes S218 and S222 to directly interact with the active site through helix unwinding, phosphorylation likely modulates the active site dynamics and plays a role in activation. In addition, as shown in Figure S1, R189 and R234 are consistently among the residues with the highest mutual information with SASA. In the double-phosphorylated state and somewhat in the S222-phosphorylated state, both R189 and R234 are attracted to phosphorylated S222 and/or phosphorylated S218, suggesting that the phosphorylated serines may be using these arginines with high mutual information to “control” the dynamics of MEK1.

We hypothesize that the DFG motif, proposed to be a conserved activation mechanism in kinases, may flip conformations in response to the presence of an ATP molecule rather than dynamical changes in the kinase itself, as in the unphosphorylated crystal structure of MEK1, the DFG motif was attracted to the ANP ligand.

Since we did not simulate MEK1 in complex with ERK2, it is still unknown how the dynamics of the critical catalytic residues contribute to the successful phosphorylation of ERK2’s phosphorylation sites. In addition, it may be worthwhile to examine the structures of the specific amino acids that have high mutual information with the SASA of the critical catalytic residues, to determine a more detailed mechanism of kinase activation by assessing the roles of individual residues.

## Methods

The PDB entry 4U7Z (MEK1 bound to G805) is selected as the protein model for molecular dynamics simulations. The missing residue 223 was built into the protein structure due to its proximity to S222, a phosphorylation site of MEK1. On the other hand, a harmonic bond restraint was placed between residues 275 and 306 to simulate a the missing intrinsically disordered loop between residues 276-305. The ligand G805 was removed from the pdb structure while ATP and the coordinating MG2+ ion are kept. To phosphorylate MEK1, the CHARMMGUI server (*31*) was used to place -2 charged phosphate group on the interest of residue(s).

The molecular dynamics simulations were run using OpenMM (*32*) on GPUv100 clusters at the University of Texas Southwestern Medical Center BioHPC computing facility. The CHARMM36 (*33*) forcefield, which contains templates for phosphorylated residues in addition to ATP, is used. The protein was first solvated in TIP3P waters (*34*) with a minimum distance of nanometer between the solute and the edge of the box. Charged ions (K and Cl) were added to the solvent to neutralize the system and to reach the physiological salt concentration of 0.15 molar. Particle mesh Ewald (PME) (*35*) is used for long-range electrostatic interactions. The temperature of the simulation is 300 Kelvin, and the pressure set to 1 bar. A timestep of 2 femtoseconds is used. All simulations were run for a total of 2 microseconds. Simulation analyses were performed using MDTraj (*36*), while mutual information analysis was performed using the ci-MIST pipeline (*28*). All visualizations were performed using VMD (*37*) and PyMol (*38*) software packages.

## Author contributions

M.M.L and E.J. conceived the work, E.J. performed the research. M.M.L and L.S. supervised the work. E.J., M.M.L, and L.S. wrote the paper.

## Supplementary Materials

Supplementary Text

## Supplementary Information

**Figure S1:**
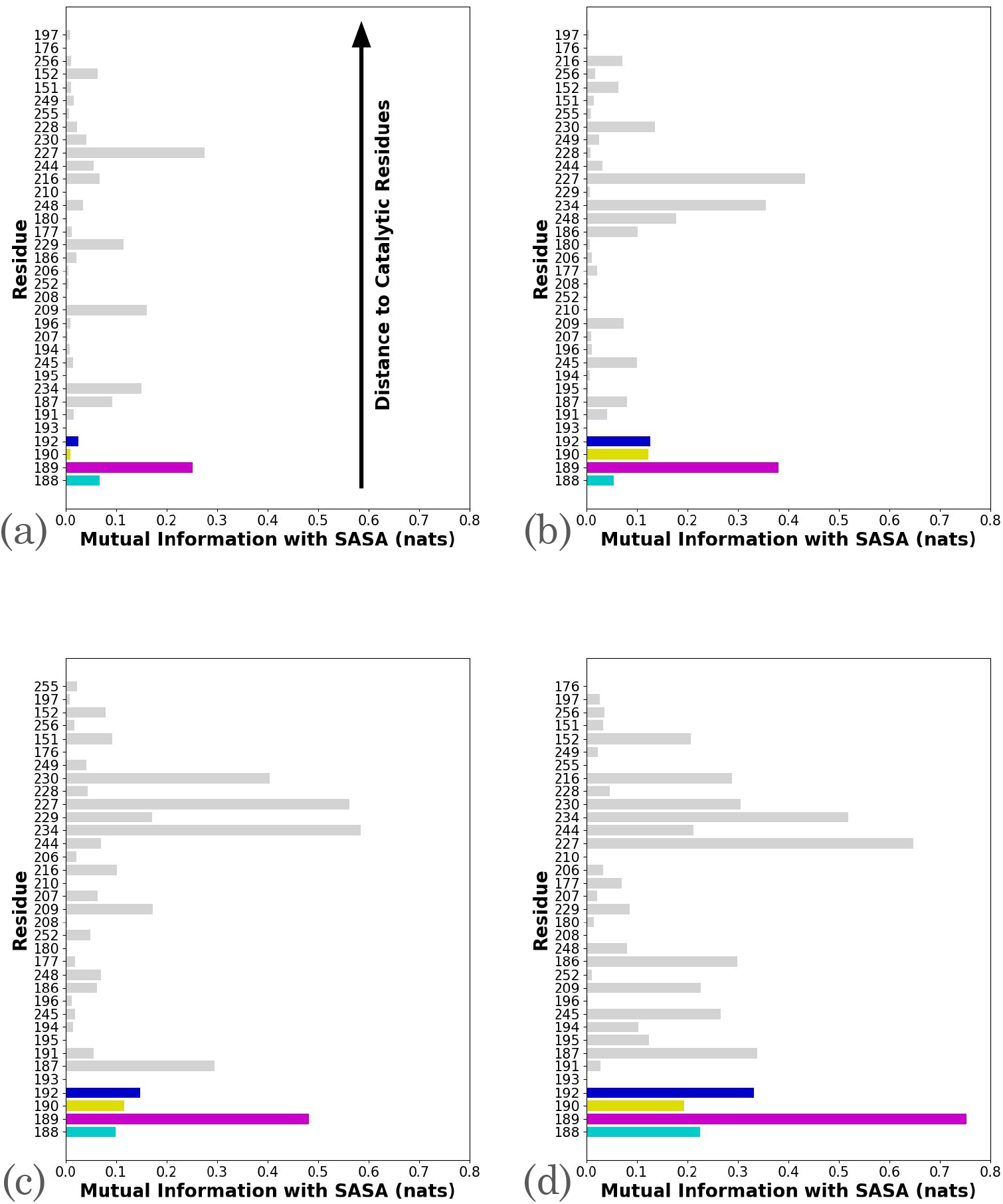
Bar graphs showing the mutual information of various residues close to the active site with SASA. The residues towards the bottom of the graphs are closer to the active site. Cyan represents H188, magenta represents R189, yellow represents D190, and blue represents K192. (a), (b), (c), and (d) represent unphosphorylated, S218-phosphorylated, S222-phosphorylated, and double phosphorylated MEK1, respectively.

